# Local and collective transitions in sparsely-interacting ecological communities

**DOI:** 10.1101/2021.10.26.465882

**Authors:** Stav Marcus, Ari M. Turner, Guy Bunin

## Abstract

Interactions in natural communities can be highly heterogeneous, with any given species interacting appreciably with only some of the others, a situation commonly represented by sparse interaction networks. We study the consequences of sparse competitive interactions, in a theoretical model of a community assembled from a species pool. We find that communities can be in a number of different regimes, depending on the interaction strength. When interactions are strong, the network of coexisting species breaks up into small subgraphs, while for weaker interactions these graphs are larger and more complex, eventually encompassing all species. This process is driven by emergence of new allowed subgraphs as interaction strength decreases, leading to sharp changes in diversity and other community properties, and at weaker interactions to two distinct collective transitions: a percolation transition, and a transition between having a unique equilibrium and having multiple alternative equilibria. Understanding community structure is thus made up of two parts: first, finding which subgraphs are allowed at a given interaction strength, and secondly, a discrete problem of matching these structures over the entire community. In a shift from the focus of many previous theories, these different regimes can be traversed by modifying the interaction strength alone, without need for heterogeneity in either interaction strengths or the number of competitors per species.

---

Interactions between species play important roles in shaping ecological communities. A central challenge in community ecology is to relate properties of interactions, such as their strength and organization, to characteristics of communities such as diversity and response to perturbations. In modeling, theory and simulations, some of the potential interactions are assumed to be negligible or irrelevant and are taken to be zero, a property known as sparseness.

Broadly speaking, theoretical approaches vary with the level of sparseness. On the sparse side of this continuum, i.e., when many of the interactions are zero, studying the structure of the network of interactions has been fruitful [1]. Many phenomena have been studied, including extended properties such as percolation, and more local properties, such as the distribution of degree (number of species interacting with each species). An extensive body of work looks at local patterns within the network [1–5] known as network modules or motifs. Central and on-going questions within this line of investigation include: whether these local patterns are more common than some null expectation; whether they play a functional role [6, 7]; whether it is possible to build-up from local properties to ecosystem-level properties such as diversity [8, 9]; and whether the ignored “weak” links can indeed be neglected [10].

In the other limit, when many or all possible interactions are present, techniques have been developed [8, 11–19] that relate the interaction strengths to properties such as the diversity, existence of multiple stable states, and persistent dynamics. Here two approaches have been used to model the community. In one, the dynamics is linearized around a fixed point, and the parameters describing the dynamics of coexisting species are sampled at random. This approach predicts stability bounds [11, 14], and has been applied to sparse interactions [20].

In the other approach, known as community assembly, the dynamics of species from a regional species pool is run, possibly resulting in the extinctions of some of the species. One interesting observation within the assembly approach, is that there are sharp transitions in many-species communities, where persistent fluctuations, very many alternative equilibria, or other properties emerge abruptly as relevant interaction characteristics are changed [8, 11–13, 15, 17, 18]. These characteristics are typically extended over the entire community (e.g., moments of interaction strengths distribution) [21]. These transitions are known as collective transitions, because they arise from community-wide processes, and a result of this is that they become sharp in the many-species limit. Whether and how these phenomena are found when interactions are very sparse (with a finite number of links per species), and whether they are at all related to local connectivity patterns that have been discussed for sparse systems, has received little attention.

Here we find that sparsely-interacting communities can exhibit phenomena associated with both lines of investigation, in different regimes, depending on interaction strength. In a theoretical model where a community is assembled from a species pool, we study equilbria and find that when interactions are strong, subgraphs of finite size play a defining role in coexistence: the problem of species coexistence reduces to a discrete problem on graphs involving local rules, in the spirit of network motifs. The coexisting species can be separated into connected sub-graphs of the interaction network see Fig. 1(A). These play a central role in our theory. The number of possible subgraph structures grows as the interaction strength is lowered, with the subgraphs typically increasing in size. The addition of each new allowed structure is marked by a transition in diversity and species abundance distributions. Note that this would not be possible in a fully-interacting community, which cannot break into multiple connected subgraphs.

**Figure 1.**
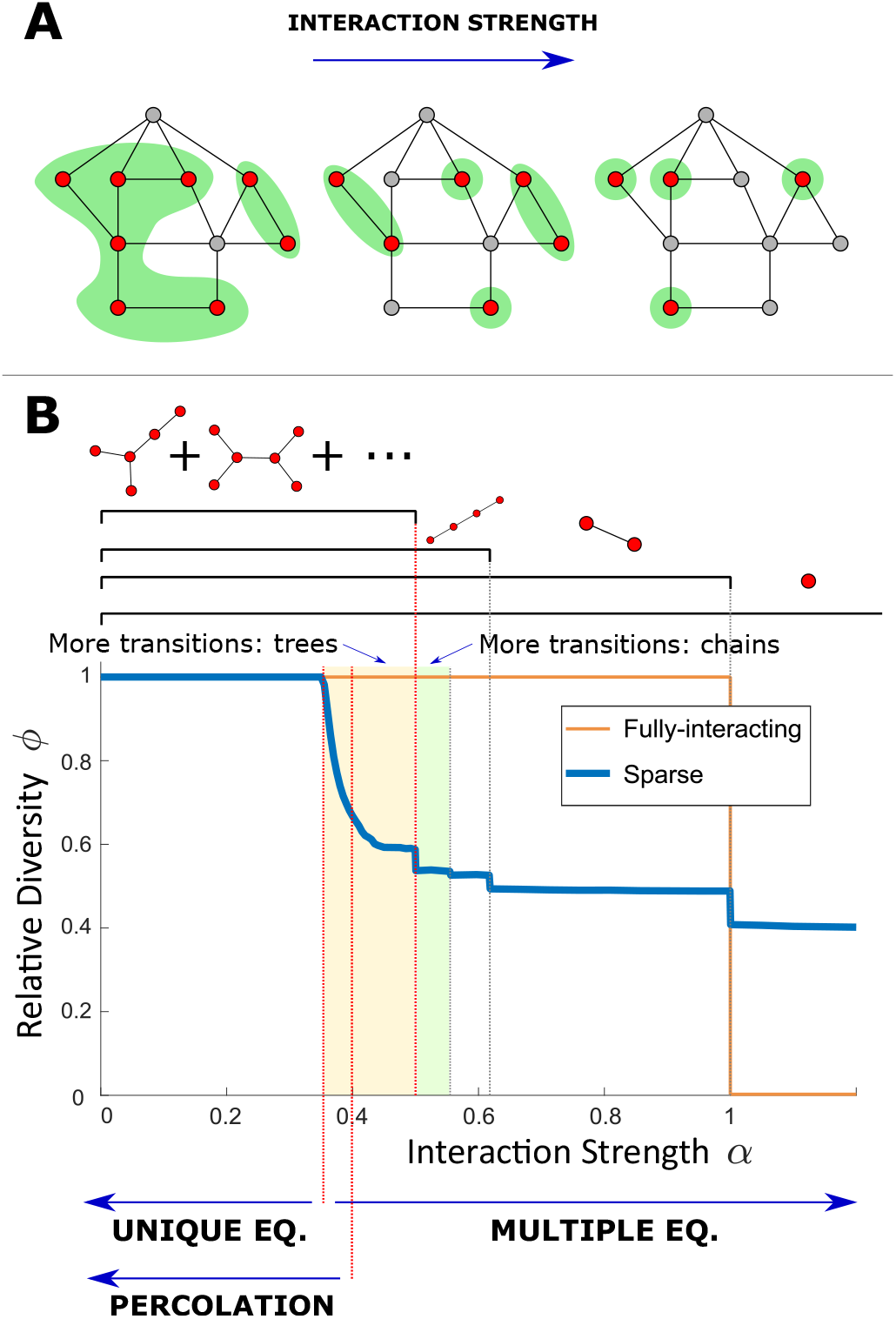
Transitions in community structure. (A) Equilibria reached at different interaction strengths. Red vertices represent persistent species, gray extinct species, and edges mark pairs of interacting species. The persistent species can be divided into connected subgraphs, shown with green background, separated by the extinct species. As the interaction strength *α* is increased, there are fewer and typically smaller allowed subgraphs, reducing the number of coexisting species. (B) The relative diversity *ϕ* = *S*\*/*S* (where *S*\* is the number of persistent species) at equilibrium, from simulations with pool size *S* = 400 and sparse interactions with degree *C* = 3 (blue); and for comparison for a fully-interacting community (orange), which exhibits only a single jump at *α* = 1. The sparse case exhibits infinitely-many sharp transitions, some of which are marked by dashed vertical lines. By order of increasing *α*, the first transition is at 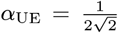, from a unique to multiple equilibria. Next there is a percolation transition at *α*_perc_ ≈ 0.4, below which a finite fraction of the persistent species belong to a single giant connected component. The other transitions result from changes in the allowed connected subgraphs specifically those that are trees (see text), leading to jumps in *ϕ*. Some of the allowed trees are shown above the graph. At values above 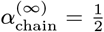, the only allowed trees are chains of even length, each with a its own allowed range. The allowed regions for trees that are not chains terminate at different values of *α* depending on the tree, all with *α* < 1/2 (shaded in orange).

At lower interaction strengths, as the interaction strength is varied we find a percolation transition, and a transition between unique and multiple alternative equilibria, similar to ones found in fully-interacting systems [22, 23].

Interestingly, all these phenomena do not require heterogeneity in either the degree or the strength of interactions. In fact, the interactions may even be locally ordered, that is, almost all species can have identical neighborhoods up to a finite distance in the network. This is in contrast to collective transitions studied previously, in which heterogeneity is necessary for the transitions to occur [8, 11–17]. The interaction strength thus becomes an important parameter on its own, divorced from the width of the distribution.

The paper is organized as follows. Sec. I introduces a theoretical model of a sparsely interacting competitive community assembled from a species pool, in which each species interacts with the same number of other species, and all interaction are of identical strength. The properties of equilibria at different interaction strengths are discussed. Interacting subgraphs of coexisting species are introduced and their role is elucidated. Jumps in diversity, a percolation and a unique- to multiple-equilibria transitions are found. Sec. II extends the model to include heterogeneity in the network of the vertex degree and interaction strengths. Sec. III shows how connected subgraphs form by combinations of smaller ones. Sec. IV concludes with a discussion.

## I. Constant Sparse Interactions

### A. The model

We work within the framework of species assembly, where species migrate from a species pool, and interact inside a community. The abundances change in time according to the standard multi-species Lotka-Volterra equations. There are *S* species in the pool. The abundance of the *i*-th species, *N*_*i*_, follows the equation

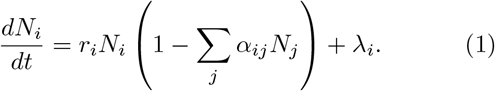

where *α*_*ij*_ are the interaction coefficients, *r*_*i*_ the growth rates, and *λ*_*i*_ the migration rates.

In this paper the matrix *α*_*ij*_, called the interaction or community matrix [24, 25], is always assumed to be symmetric, *α*_*ij*_ = *α*_*ji*_, with equal intraspecific competition for all species, *α*_*ii*_ = 1. The symmetry ensures that the dynamics in Eq. (1) always reaches an equilibrium [26]; There may be one or more such equilibria. Here we only consider competitive interactions, *α*_*ij*_ ≥ 0, and assume that all growth rates are positive, *r*_*i*_ > 0; other than that the values of the *r*_*i*_’s have no effect on the set of stable equilibria. In simulations we take all *r*_*i*_ = 1, and run Eq. (1) until changes in the *N*_*i*_-values are small. The migration strengths *λ*_*i*_ are taken to be small, *λ*_*i*_ → 0^+^, ensuring that at an equilibrium (i.e. a stable fixed point), all species that could invade do so. We use a migration rate of *λ*_*i*_ = 10^−10^, and species are considered extinct when *N*_*i*_ < 10^−5^. To ensure a true equilibrium has been reached, it is verified that 1 – Σ_*j*_ *α*_*ij*_*N*_*j*_ = 0 for all present species with *N*_*i*_ > 0, and that extinct species cannot invade, *dN*_*i*_/*dt* < 0.

We are interested here in sparse interactions, where many of the pairs of species do not interact (*α*_*ij*_ = 0). The network of interactions forms an undirected graph, with vertices representing species and edges representing pairs of interacting species, sometimes called the community graph [27].

It is common to use random interactions sampled from different distributions, which capture different interaction characteristics. In this section we will consider the following model: (1) Each species interacts with exactly *C* other species, with the interacting pairs chosen at random so that the community graph is a random *C*-regular undirected graph. (2) The interaction strength is equal for all interacting pairs. Therefore, the interaction matrix can be written as *α*_*ij*_ = *δ*_*ij*_ + *αA*_*ij*_, where *A*_*ij*_ is the symmetric adjacency matrix of the community graph. We consider *C* ≪ *S*, and more precisely the limit of large *S* at constant *C*. We will see that this simplified model already yields dramatically different results as compared with the fully-connected system. Extensions to varying interaction strength and number of interaction per species are then discussed in Section II.

We limit the discussion to properties of the system’s equilibria, and not the dynamics towards the equilibria, or under additional noise, which are very interesting (some already discussed in [28]) but beyond the scope of this work.

### B. Overview of different regimes

To get a bird’s eye view of the different behaviors, we follow the diversity at the equilibria as a function of the interaction strength α (recall that in this first model α is identical for all pairs). Let *S*\* be the number of co-existing species at an equilibrium (species richness), and define the relative diversity *ϕ* = *S*\*/*S*, their fraction relative to the total number *S* of species in the pool. Fig. 1 shows simulation results for *ϕ* as a function of *α*. *ϕ* is estimated by running simulations of Eq. (1) over many realizations of adjacency matrices *A*_*ij*_, starting from a few different initial conditions per realization, with each *N*_*i*_ sampled uniformly from [0, 1]. The variability in *ϕ* between simulations under the same conditions decreases with the diversity *S*, and for large *S* it is essentially set deterministically. For comparison, the case of a full community matrix, where all species interact with each other with strength *α* is also plotted. In this case the behavior is simple: For *α* < 1 there is a unique fixed point in which all species are persistent and *ϕ* = 1, while for α > 1 there are S different fixed points, each with a single persistent species so that *ϕ* = 1/*S*, tending to zero at large *S*. The sparsely-interacting system, in contrast, is very rich and exhibits multiple different behaviors with sharp transitions between them. At values of α close to zero, the interactions are weak enough to allow all species to coexist with *ϕ* = 1. This persists for larger *α* up to some critical value *α*_UE_ where *ϕ* starts to decrease. At another value *α*_perc_ there is a percolation transition, above which none of the components of persistent species scales with the system size. The relative diversity *ϕ* keeps decreasing until it reaches another transition where there is a jump in *ϕ*, at a value we denote by 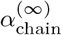. At 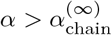, the relative diversity *ϕ*(*α*) consists of infinitely many plateaus punctuated by jumps, until the last jump at *α* = 1 and a single plateau above it.

In the following sections, we discuss this behavior in detail, and explain the multiple changes in system behavior and the reasons behind them. We will show in the next sections that 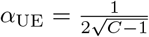 and 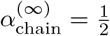, and provide analytical values for *α* of all jumps in *ϕ*(*α*) at *α* ≥ 1/2. In Subsection I D we discuss the percolation transition, and in Subsection I E the unique to multiple equilibria transition, and show that it coincides with *α* where *ϕ* first drops below 1.

### C. Allowed subgraphs and their dependence on interaction strength

Here we begin to explain the different regimes described in Section I B, by analyzing properties of the equilibria of the model. In the limit of small migration (*λ*_*i*_ → 0^+^) some of the species will persist (*N*_*i*_ > 0) and others go locally extinct (*N*_*i*_ = 0 as *λ*_*i*_ → 0^+^). At an equilibrium, the extinct species must be unable to invade (*dN*_*i*_/*dt* < 0), and the abundances of the persistent species must return to the fixed point if perturbed away from it. These conditions will be referred to as *uninvadability* and *stability*, respectively. The persistent species can be grouped into connected subgraphs of the community graph, see Fig. 1(A).

We begin in the limit of very large *α*, studied in [8, 29]. Under this very strong competition, the problem reduces to two conditions. First, two interacting species cannot both persist (competitive exclusion). The connected subgraphs are thus individual species, see Fig. 1(A), rightmost illustration. Second, an extinct species cannot invade if and only if it interacts with one or more persistent species. Stability is automatically satisfied, as it involves isolated persistent species. Importantly, the values of *α* do not appear in these two conditions, and so finding an equilibrium point reduces to a discrete, combinatorial problem on the graph, of finding a maximally independent set [8]. In [29], the authors used this insight to calculate the diversity and number of equilibria on Erdős–Rényi graphs (where the pairs of interacting species are chosen independently with some probability).

At lower values of *α* the connected subgraphs are no longer only isolated species, see Fig. 1(A). These subgraphs must satisfy certain “internal properties” in order for them to appear at a given *α*. As long as all of the neighboring species to the subgraph are extinct, the abundances at a fixed point of the subgraph are determined entirely by interactions within it. These abundances must be positive (a condition known as *feasibility*), and the fixed point must be stable. These conditions depend only on *α*. This allows us to understand much of the behavior by looking at individual subgraphs: each subgraph *μ* will have a critical value 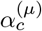, above which it is either unstable or not feasible, and can therefore only appear at an equilibrium of a system in the “allowed” range 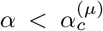. (This leaves out a possibility that a graph could switch back and forth between being allowed or not, see Appendix A.) Thus the system is governed by discrete combinatorial conditions, which determine the entire set of possible equilibria of a given system.

Here another important simplification enters. Sparse random graphs, including random-regular graphs and Erdős–Rényi graphs discussed in Sec. II, are locally tree-like, meaning that they have only a finite number of short cycles even when *S* is large. For example, in a large random regular graph with *C* = 3 the average number of triangles is 4/3 [30]. Thus, most connected subgraphs of finite size in the network will be trees, i.e., contain no cycles, and properties such as diversity and species abundance distribution that are averages over the entire community can be calculated by only considering trees, and specifically, the critical values 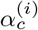 need to be found only for trees. Examples of connected subgraphs within a local tree neighborhood are shown in Fig. 6 in Appendix A.

The trees can be divided into chains and other trees. We calculated 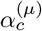 for chains analytically, see Appendix A. For a chain with *n* species,

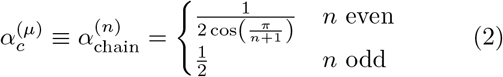

For chains of even length, 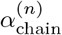 is a decreasing series that converges from above to 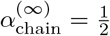, which is also the critical value for all chains of odd length, 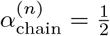. All other trees have 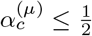, with the first ones appearing, coincidentally, exactly at 1/2, as we prove in Appendix A. Therefore, 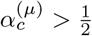 only for chains of even length, so only they can appear in communities at *α* > 1/2.

In addition to these “intrinsic” considerations about the stability and feasibility of different connected components, uninvadability must also be considered. This is more complex since it depends on how the components fit together, and in principle this could lead to additional jumps in *ϕ*, but in the *α* > 1/2 region, such jumps seem to be rare if they exist at all, and their size is so small that we have not detected them in simulations. See details in Appendix B.

This means that for *α* > 1/2, in ranges of *α* between the 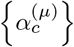, the same trees will be allowed and so essentially the same set of equilibria will exist (since uninvadability does not seem to be important except at the transitions). As *α* is lowered below some 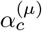, a new tree abruptly appears, leading to many new possible configurations and thus causing the diversity to jump. While the dynamical simulations used to obtain *ϕ* (*α*) do not necessarily reach all equilibria with the same probability, they clearly show jumps in *ϕ* at these values, with plateaus of approximately constant values of *ϕ* in between. Fig. 1(B) shows the function *ϕ* (*α*), marking some of the critical 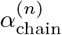 from Eq. (2) as dashed vertical lines, showing that the jumps in *ϕ* indeed happen exactly at 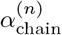. This also happens for trees that are not chains when *α* < 1/2, see an example in Appendix A.

As *α* is lowered, infinitely many subgraphs of more complex structures become stable, so the values 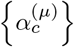 become more dense, and the jumps in *ϕ* (*α*) smaller (see Fig. 7). This makes it harder to observe them in numerics, but we expect that they exist in the entire range down to *α*_UE_, defined in the following. Once trees appear there are many interesting types of transitions that could happen. Just as at 1/2 arbitrarily long chains appear, there could be other points where there are qualitative changes in the properties of trees; see the Discussion section for more discussion.

The transitions are also reflected in the possible abundances of species, as seen in rank-abundance curves, which show the abundances sorted in decreasing order, see Fig. 2. At a given α, the abundance of a species depends only on the connected tree it belongs to, and its position within it; for example, species that belong to a chain of length two have 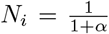. Therefore, as a tree *μ* becomes feasible and stable at 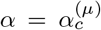, the abundances associated with it can appear at an equilibrium. As shown in Fig. 2(A), this causes the abundance graphs to smooth out as *α* is lowered, since the number of possible abundances increases.

**Figure 2.**
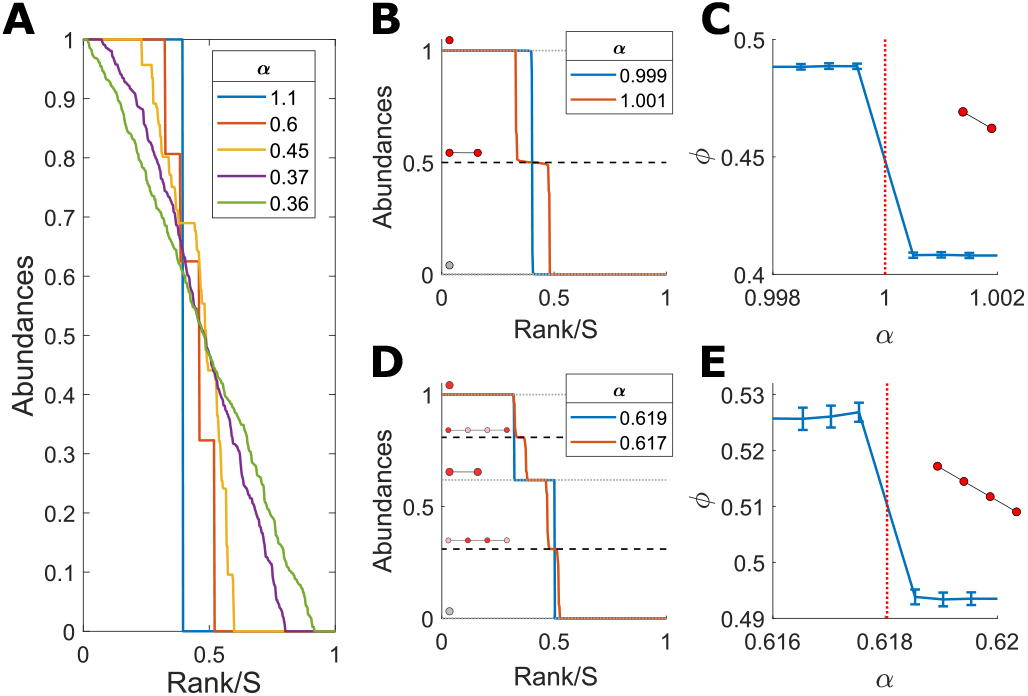
Changes in feasible and stable trees are reflected in species abundances. At each value of the interaction strength *α*, certain trees are allowed, and the abundance of a species depends only on *α* and the position within a tree. **(A)** The rank-abundance curves at equilibria reached dynamically for *S* = 400,*C* = 3, at several values of *α*. As *α* decreases, the increasing number of feasible and stable trees generates more possible species abundances. **(B-E)** Species abundances on both sides of two transitions at 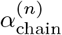 for *n* = 2, 4, where new trees appear. **(C)** and **(E)** show the behavior of *ϕ* around the transitions associated with pairs of species and chains of length 4 respectively becoming feasible and stable. **(B)** and (**D)** show the abundances at equilibria at values of *α* on two sides of the transitions. The expected abundances are marked by dashed black lines, with thicker lines for the abundances of the species in the tree associated with the transition. Next to each abundance appears the tree that contains it, with the species that have this abundance in dark red (or gray in the case of the abundance 0 of extinct species).

To summarize, in this section we described how the interaction network breaks up into connected subgraphs, with changes in allowed subgraphs driving jumps in diversity and species abundances. These subgraphs are trees that are feasible and stable at that interaction strength. Finding the equilibria of Eq. (1) reduces into a discrete graph theoretical problem on the community graph. Broadly speaking, for stronger competition there are fewer and typically smaller allowed trees.

As *α* is lowered, the size of the allowed subgraphs grows until they span a finite fraction of the species, as discussed in the next section. The number of different types of allowed graphs quickly grows with their size, and the problem of classifying them becomes more difficult, and less useful. These very large connected graphs can include the rare but still existing cycles in the graphs, and so they are no longer trees.

### D. Percolation transition

Percolation transitions are one of the canonical phenomena studied in graph theory. In site percolation, some vertices of a graph are removed. As the probability of vertex removal varies, on one side of the transition the remaining graph breaks into small (sub-extensive) pieces; on the other side, a finite fraction of vertices belong to a single connected component. Natural communities belonging to both regimes are known to exist [1].

We find that at some interaction strength *α*_perc_ there is a percolation transition, below which the largest connected subgraph formed by surviving species is extensive, that is, includes a finite fraction of all the species. Fig. 3(B) shows the fraction of species belonging to the largest connected component as a function of *α*, for several values *S* with *C* = 3. Above a certain *α*, which for this connectivity is at *α*_perc_ (*C* = 3) ≈ 0.41 ± 0.01 (marked by a dashed line), this fraction drops with *S* indicating a sub-extensive largest component. Below *α*_perc_ this fraction converges to a constant value. As expected, this value is smaller than 1/2, since at 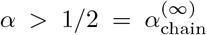 the only possible components are finite-length chains, as shown in Sec. I C above. Also, *α*_perc_ ≥ *α*_UE_ where all species persist, see Sec. I E below. The fact that the transition becomes sharper with growing S is a hallmark of a collective transition.

Fig. 3(B) is qualitatively similar to that of a standard site-percolation transition, where vertices are randomly and independently chosen to be “present”, see Appendix D. However, the fraction of persistent species at *α*_perc_ is around *ϕ*_perc_ ≈ 0.64 ± 0.02, which is larger than the *ϕ*_perc_ = 1/2 of a standard site percolation transition at *C* = 3 [31]. This is because in our model, the species that persist are not sampled independently; the higher value in our model is expected given that persistent species are correlated, tending not to be adjacent to one another.

**Figure 3.**
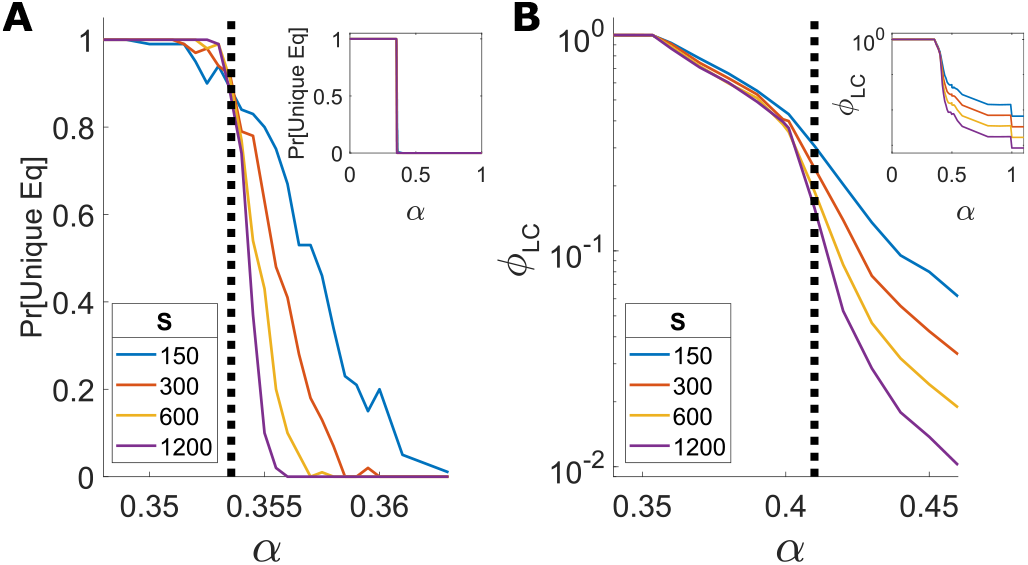
Collective transitions. **(A)** Unique to multiple equilibria transition: the probability for a unique equilibrium as a function of *α*, for connectivity *C* = 3 and several pool sizes *S*. The probability is obtained by generating many realizations of interaction matrices and determining whether there is a unique equilibrium by the stability of the fully-feasible fixed point, as described in the text body. The exact value for the transition is shown as a dashed black line, Eq. (3). Inset: the same graph over a larger range of *α*. The transition becomes sharper with *S* grows, as expected from a colective transition. **(B)** Percolation transition: the fraction of species in the largest connected component as a function of *α*, for several values of *S*. The location of the transition is *α*_perc_ ≈ 0.41 ± 0.01 (dashed black line). At *α* < *α*_perc_ a finite fraction of species belongs to the largest component even when *S* grows. At *α* > *α*_perc_, this fraction decreases with *S*. Inset: the same graph over a larger range of *α*. Here too, the transition becomes sharper at larger values of *S*.

### E. Unique to multiple equilibria transition

The final transition in the model with all-equal *α*, at the lowest value of *α*, is from multiple to unique equilibria. In order to find the critical value of *α* for this transition, we first argue that the community has a unique equilibrium exactly when it is “fully feasible”, i.e. all species are persistent (*ϕ* = 1); if the fully-feasible state is an equilibrium then it is necessarily unique. Thus, the transition from the multiple equilibria phases to the unique equilibrium phase occurs at the value *α*_UE_ (*C*) in which *ϕ* drops below 1. The equivalence holds only for this model where all species have the same number of interacting pairs and all interactions have the same strength *α*, and breaks in more general cases, see Sec. II below.

To understand this relation, consider the Lyapunov function *F* = 2 Σ_*i*_ *N*_*i*_ – Σ_*ij*_ *N*_*i*_*α*_*ij*_*N*_*i*_, which for symmetric interactions (*α*_*ij*_ = *α*_*ji*_) grows with time according to the Lotka-Volterra equations [26], and whose local maxima coincide with the equilibria. The fixed point where all species persist is always feasible, as from the local homogeneity of the community graph all abundances are equal 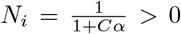, and this would be stable if the full interaction *α*_*ij*_ is positive definite. As *α*_*ij*_ is also the matrix of second derivatives of *F*, if the fixed point is stable then the Lyapunov function is concave everywhere, meaning the minimum at the “fully feasible” equilibrium is global and therefore unique.

Conversely, if the fully feasible equilibrium is not stable, then *F* is a non-concave quadratic function on the quadrant {*∀i*: *N*_*i*_ ≥ 0} and one expectsthat if there are many potential species, it is likely to have many local maxima, and therefore multiple equilibriaWe checked this relation numerically, by generating 100 realizations of the interaction matrix at a given *α*, solving the dynamics in Eq. (1) with 30 different randomly chosen initial conditions, and checking whether all runs converge to the same equilibrium. This process was repeated around the transition (whose position is given below in Eq. (3)), for *α* ∈ [0.35, 0.36] when *C* = 3 and for *α* ∈ [0.285, 0.3] when *C* = 4, and with *S* = 200, 400. In all runs, there was a unique fixed equilibrium at exactly the same realizations that were fully feasible.

The stability is thus determined by the range in which the matrix *α*_*ij*_ = *δ*_*ij*_ + *αA*_*ij*_ is positive definite. *A*_*ij*_ is an adjacency matrix of a *C*-regular graph of size *S*, and at large *S* its minimal eigenvalue is with probability one at 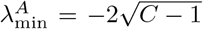 [32]. The minimal eigenvalue of the matrix *α*_*ij*_ is therefore at 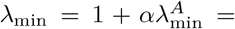 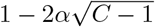, and the critical value of *α* will be

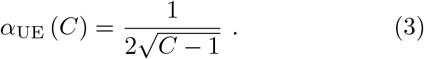

Fig. 3(A) shows the probability of the system having a unique equilibrium as a function of *α* for several values of *S*, using the stability of the matrix *α*_*ij*_. As *S* increases, the probability for a unique equilibrium becomes sharper (again, a clear sign of a collective transition), approaching a step function at the expected value of the transition *α*_UE_ (C).

## II. Heterogeneity In Vertex Degree And Interaction Strength

So far, Sec. I analyzed a model where each species interacts with exactly C others, and all with the same interaction strength α. Here we consider the effects of heterogeneity, both in the strength of species interactions and in the vertex degree (the number of species interacting with a given one). The interaction strength is varied by drawing it from a normal distributions with mean *α* and a given standard deviation *σ*. The degree is varied by replacing the random regular graphs with an Erdős-Rényi random graph, in which each pair of species is independently chosen to interact, such that the average degree is *C*. To understand how these two changes affect the results, we consider them separately. Fig. 4 shows the relative diversity *ϕ* as a function of *α*, for both cases.

**Figure 4.**
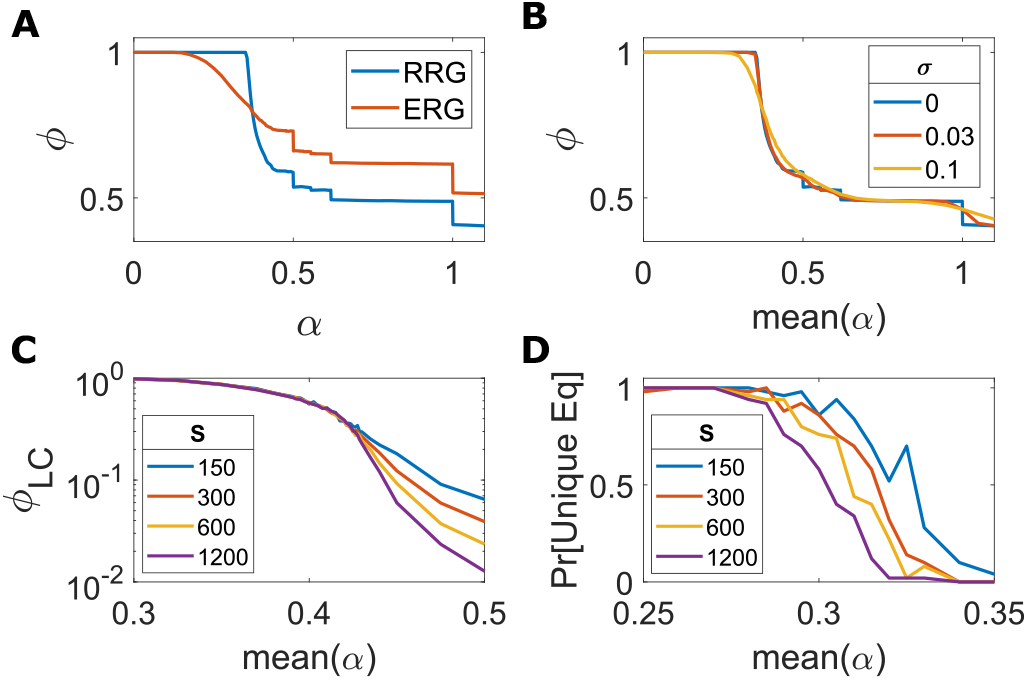
Transitions with heterogeneous interaction strengths and degrees. **(A-B)** Transitions due to changes in allowed trees are broadened when there is variability in interaction strengths, but remain sharp for variation in degree. The relative diversity *ϕ* as a function of interaction strength *α*, for *S* = 400, *C* = 3. **(A)** Erdős-Rényi graphs with interaction probability *p* = *C/S*, compared to a random *C*-regular graph. **(B)** Interaction strength is drawn from a normal distribution with mean *α* and standard-deviation *σ* = 0, 0.03, 0.1, keeping the interactions symmetric and a random regular graph. **(C-D**) The collective transitions with heterogeneity in interaction strength become sharper as *S* increases, just as they do without it (compare with Fig. 3). Results are shown for *C* = 3, *σ* = 0.1 and several values of *S*. **(C)** Percolation transition: The fraction of species in the largest connected component as a function of *α*. **(D)** Unique to multiple equilibria transition: The probability of having a unique equilibrium as a function of *α*.

When varying the degree, the jumps in the relative diversity *ϕ* due to changes in the allowed trees remain sharp, while they are broadened for variations in interactions strength. This makes sense, as the trees can still exist if the degrees vary; there may be additional adjacent species but these do not affect whether the populations on the tree are feasible and stable. On the other hand, the interaction strengths affect the stability and feasibility of the tree. In an Erdős-Rényi graph, all trees of the same topology will all have the same *α*_*c*_. (*ϕ* is different between the Erdős-Rényi and regular graphs due to their different structure.)

If interaction strengths are varied, trees in the same system, which have the same topology but different interaction strengths might have different limits on stability and feasibility, leading to the appearance of more types of allowed trees than in the all-equal *α* case. But if the disorder is not too strong, the picture of the all-equal *α* case remains relevant: if the mean value of *α* is within a plateau of the all-equal *α* case and not too close to the ends, the interactions would mostly allow the same trees as they would in the case without disorder. For example, for mean (*α*) = 0.7, within the plateau allowing only pairs and singlets, for *σ* = 0.1 these make up 99.5% of feasible trees in a typical equilibrium for large S. Indeed, for *σ* = 0.1, in most of the range within this plateau, *ϕ* is almost identical to the all-equal *α* case Fig. 4(B).

The two remaining transitions, for percolation and from unique to multiple equilibria, both appear to become sharper as S increases for both variations in interaction strengths and degree, as can be seen in Fig. 4(C-D) and in Appendix C, Fig. 10. For any given *S* the transitions are broader compared to the equal-*α* model (Fig. 3). Furthermore, in both cases *ϕ* drops below 1 while still at the unique equilibrium phase, which happens when the system is no longer fully feasible. This is clearly seen for the Erdős-Rényi random graph in Fig. 4(A), and is shown for varying interaction strengths in Appendix C. This is in contrast to the all-equal *α* model, where *ϕ* drops below 1 when the system becomes unstable at *α*_UE_.

## III. Subgraph Emergence Rule: How The Trees Grow

As the interaction strength is lowered (by lowering *α* in Sec. I A or mean (*α*) in Sec. II), the allowed connected subgraphs become larger (containing more species) and more complicated, until one connected subgraph can take up a finite fraction of the community at the percolation transition. Continuing to grow beyond that, they finally include the entire network. For *α* > 1/2 there is a clear regularity in the sequence of transitions, as even-length chains become allowed by order of length. This raises the question of whether there is any regularity by which more complicated subgraphs (trees, and even subgraphs with cycles) become allowed. We now describe a general and simple result, when the interactions strengths are heterogeneous.

Consider a subgraph within the interaction network, see Fig. 5. Since interaction strengths are not all equal, this refers to a specific set of vertices, which means the result can be different for the same subgraph structure when it involves different species. To define *α*_*c*_ of the sub-graph, consider the process by which the mean strength is changed by shifting the values of the *α*_*ij*_, i.e. adding a constant (other continuous changes of the matrix *α* are also possible). Just below *α*_*c*_ all abundances are positive. We prove in Appendix E that with probability one, it is feasibility, rather than stability, that is lost at *α*_*c*_, by one species going extinct (*N*_*i*_ → 0). When this becomes extinct, the remaining subgraph is composed of allowed subgraphs, hence the subgraph right below *α*_*c*_ is thus composed of subgraphs allowed right above *α*_*c*_, with one additional vertex.

**Figure 5.**
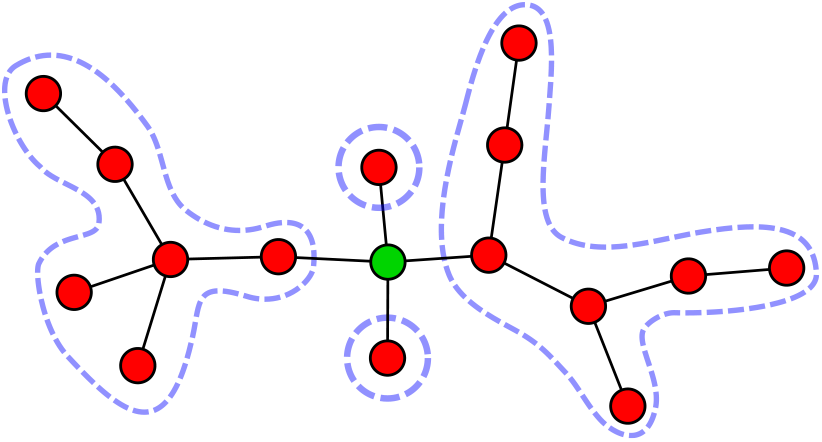
A tree that becomes feasible and stable at some *α*_*c*_, can be constructed from three or more trees that are allowed right above *α*_*c*_ (surrounded by dashed lines), joined by one additional species (green). This is true with probability one when there is heterogeneity in interaction strengths.

For a tree subgraph, the species that goes extinct interacts with three or more species in that subgraph (is a branching point), assuming that the distribution of the *α*_*ij*_-values is not too wide. See argument in Appendix E. This implies that the tree splits into three or more trees.

This construction gives a constraint on what order the specific subgraphs become allowed, i.e. become feasible and stable: that when a subgraph becomes allowed as *α* is decreased, pieces of it with one species removed were already allowed. As noted above, because of the heterogeneity of the interactions, here a subgraph refers to a specific set of species on which it resides. Note that as always, whether an allowed graph appears in an equilibrium depends also on the neighboring species and the rest of the network.

## IV. Discussion

We have looked at a community assembled from a pool of sparsely-interacting species. When the interactions are strong enough, the assembly process breaks the network into many connected subgraphs. The problem of equilibrium coexistence reduces to understanding which subgraphs are allowed, and how they are organized to keep extinct species from invading.

When these subgraphs are small, it might be possible to formulate predictive local rules about their occurrence, in the spirit of “assembly rules” [4, 33, 34]. The simplest example is competitive exclusion, where if the interactions between two species are greater than one, *α*_*ij*_, *α*_*ji*_ > 1, then they cannot coexist within a community of species that interact competitively, irrespective of the state of the other species. This can be interpreted as a rule that when interaction are stronger than one, the connected components include just one species. Here this regularity extends to weaker interaction strengths, first identifying a regime where interacting pairs are also allowed, which is quite robust to some level of heterogeneity in interactions strengths (Sec. II), and then to regimes with larger connected subgraphs.

For lower interaction strengths there are many larger allowed subgraphs, making the corresponding graph-theoretical problem hard and far less local, and limiting the potential for predictive local rules. There is also more sensitivity to heterogeneity in the interaction strengths. At even lower interaction strengths, connections percolate across the entire network of coexisting species, and below that there is a dramatic transition in behavior, as the equilibrium becomes unique, similar to transitions found in fully-interacting systems [12, 22, 23]. We have not observed any sharp changes occurring at the percolation transition, to diversity, stability or other measures beyond the graph connectivity; percolation might however be a necessary bridge between the finite-subgraph regime, and the unique to multiple-equilibria transition.

In a striking difference from fully-connected networks, this rich phenomenology does not require heterogeneity in interaction strengths or vertex degrees, which are necessary in fully-interacting networks, and have been central to much of the field for decades [8, 11–17]. This makes the interaction strength (or its mean as opposed to the width of the distribution) a parameter of independent importance, single-handedly driving changes in stability and feasibility. When heterogeneity is present, the mean and distribution of interaction strengths have a combined effect, with the allowed subgraphs still playing a central role in shaping the community.

There are many mathematical questions to explore in these systems, which are interesting because of the interplay between the combinatorial structure of the community graphs and the quantitative properties of the interaction matrices. Such questions include a further understanding of the sequence of transitions: Are there other limit points where infinite trees become stable (such as the infinite chain becoming stable at *α* = 1/2)? And are there ranges where the critical points 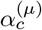 are dense? Another question is how much the dynamics affects the distribution of equilibria reached, as compared to an unbiased choice among all the allowed equilibria; for example, how much this affects the sizes of the jumps in diversity. Finally, it would be especially interesting to understand the transition to the unique equilibrium state, by studying the structure of the small groups of species that go extinct just above the transition. It would likely be possible to make progress on many of these questions by studying ideal infinite trees with a fixed degree *C*, so that the inhomogeneity in the equilibria arises from their instabilities.

The extent to which interactions in different natural communities are sparse is an open question, since directly measuring interaction strengths can be hard, especially the weaker ones. This is complicated by additional factors, as many weak interactions might have a large cumulative effect, and that some inference techniques assume that the network is sparse (e.g., [35]). One can hope that studying consequences of sparsity would help identify and better understand such communities.

## ACKNOWLEDGMENTS

It is a pleasure to thank M. Barbier, J. Friedman and J. Gore, for stimulating discussions and helpful feedback. G. Bunin was supported by the Israel Science Foundation (ISF) Grant No. 773/18.

## Appendix A Critical values of trees

The critical *α* values for connected trees, 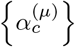, are discussed in section I C of the main text. They are the values above which each tree becomes either unstable or unfeasible, and therefore cannot appear in an equilibrium. In rare cases, a stable tree can regain unfeasibility after losing it (see below); here we define 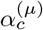 more precisely as the *highest* value of *α* where the tree is feasible and stable. As discussed in the main text, trees are important subgraphs because the local neighborhoods of most species are locally tree-like. Examples of trees appearing in an equilibrium in the neighborhood of one species within a large community are shown in Fig. 6

**Figure 6.**
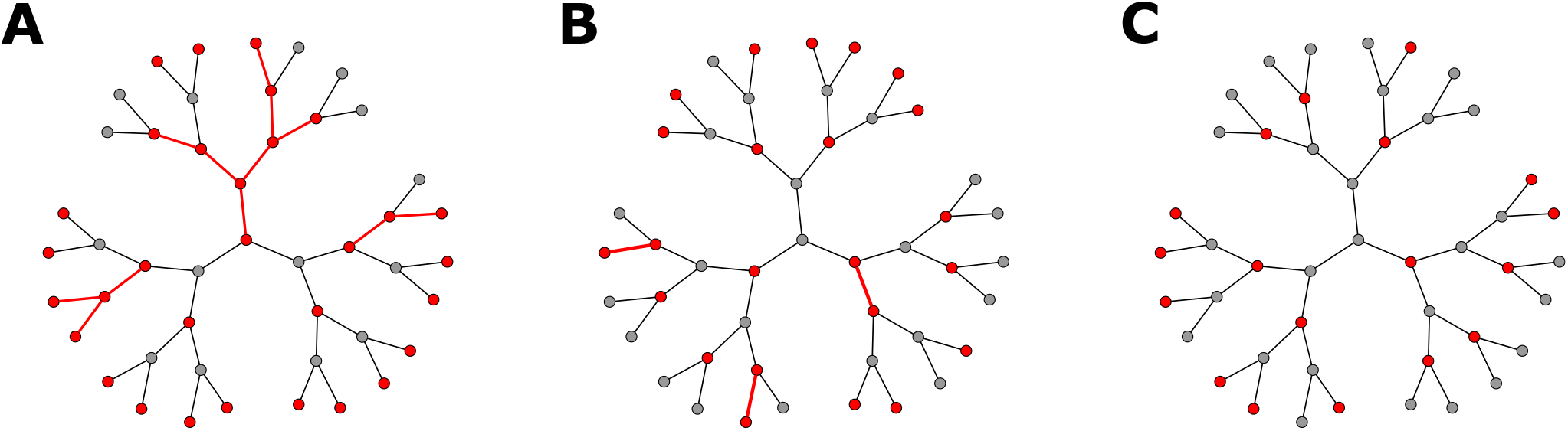
Equilibrium in the local neighborhood of one species. The neighborhood of one species in a specific system with connectivity *C* = 3 and pool size *S* = 1200, where the interaction strengths are **(A)** *α* = 0.45 **(B)** *α* = 0.7 **(C)** *α* = 1.1. Extinct species are in gray, and persistent species in red. The edges connecting two persistent species are also marked in red. As the interaction strength is increased, the subgraphs change from large trees that are not chains in (A), to length-2 chains and singlets in (B), and singlets only in (C).

Stability changes at a single value of α, so that the system is stable for all values of α below it and unstable for all values above it. Indeed, as the interaction parameters, in matrix form, are represented by *I* + *αA*, where *I* is the identity matrix (see Sec. I A), if the minimal eigenvalue of the adjacency matrix A representing a tree is *λ*_*min*_, the smallest eigenvalue of the tree at the interaction strength *α* would be 1 + *αλ*_*min*_, so the tree is unstable exactly for *α* > −1/*λ*_*min*_. Feasibility on the other hand can be gained and then lost more than once, but we find that most stable trees that gain feasibility as *α* is lowered usually retain it, with feasibility gained and lost again in only 0.03% of trees up to 20 vertices for *C* = 3 and 0.01% of trees up to 15 vertices for *C* = 5.

As mentioned in section I C, we find that for a chain of length *n*,

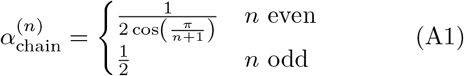

and for all other trees, 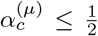, with *μ* going over all trees. A histogram of the values for small trees, calculated numerically, appear in Fig 7(A), and the example in Fig. 7(B) shows that they do indeed generate jumps in *ϕ* even in the *α* < 1/2 region.

**Figure 7.**
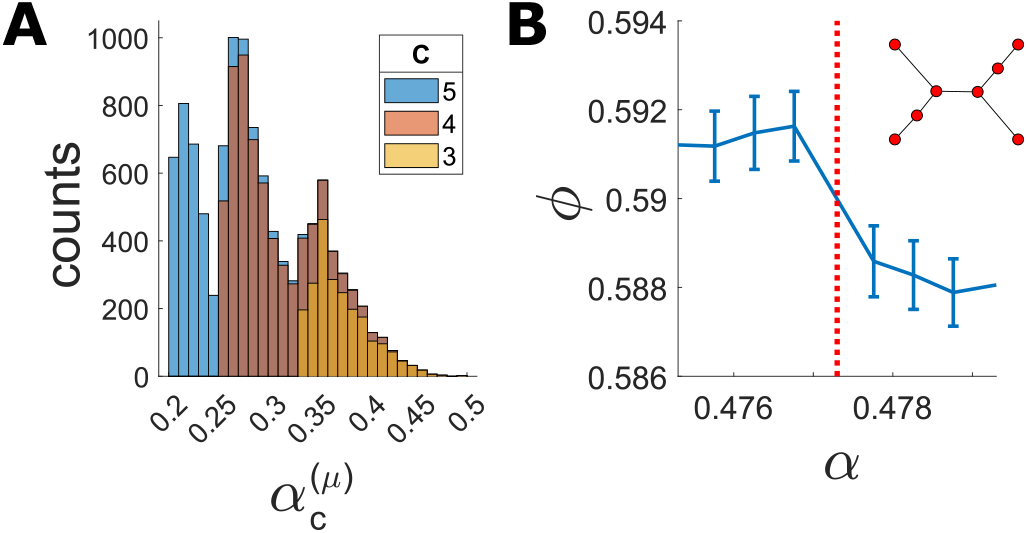
The critical values 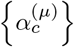 for trees that are not chains. **(A)** Histogram of the critical values for trees up to size 15, excluding chains, which are possible subgraphs of random regular graphs with connectivity *C* = 3, 4, 5. **(B)** Trees can generate jumps in *ϕ* at *α* < 1/2: behavior of *ϕ* for *S* = 400, *C* = 3 around the value 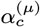 associated with the tree shown.

While it is fairly easy to determine the critical *α*’s for chains by checking stability and feasibility directly, it is hard to show that all trees that are not chains are either unfeasible or unstable above 1/2. E.g., to show they are unfeasible, we have to find the solutions to the equilibrium equations and check that one of the species has a negative abundance, but there is no general formula for inverting the interaction matrix on an arbitrary tree. Instead of doing this, we can note that if the tree is unfeasible or unstable, it will still have a stable equilibrium (because there is a Lyapunov function so it cannot oscillate indefinitely), but this stable equilibrium has missing species. It must decompose into subgraphs that are stable and feasible, which are expected to be only chains above 1/2. Conversely, the reasoning from sec I E shows that if such an uninvadable equilibrium made up of chains exists, the full graph cannot be stable and feasible. This does not require inverting the matrices for trees, just doing a more combinatorial problem of splitting the tree into pieces. This idea and another lemma are the basis for the proof.

These two lemmas, proved in Sec. A 1, are (1) A tree with an unstable sub-tree is itself unstable, and (2) Any subgraph is feasible and stable if and only if there is no stable and uninvadable equilibrium on the tree where some of the species are extinct. We then prove the results on chains, Eq. (A1) in Sec. A 2. Although this can be done directly, the lemmas give more intuitive derivations of parts of the results. For example, Lemma 2 allows us to show that an odd length chain is never feasible and stable above 1/2 by finding an equilibrium with extinct species in that range. We prove in Sec. A 3 that for trees that are not chains 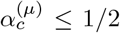, in two stages: first, we use lemma 1 to show that unless the junctions in the tree have at least one neighbor of degree 1 and the rest of degree 2 at most, the tree is unstable. Next we prove that, for any junction with these properties, an equilibrium exists for *α* > 1/2 in which the species at the junction is extinct, so by lemma 2, the tree is not feasible and stable.

## 1. Supporting Lemmas

Here we will introduce two lemmas to aid the proof. They are also of interest in their own right.

### (1) Lemma

A graph that has an unstable subgraph is itself unstable.

Proof: To prove this, denote the interaction matrices of the entire graph and the subgraph as *α*_*ij*_ and 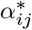 respectively, and their minimal eigenvalues as 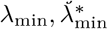, with 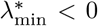 as the subgraph is unstable. As 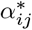 is a principal submatrix of *α*_*ij*_, from the Cauchy eigenvalue interlacing inequality 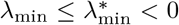, meaning the full graph is unstable.

### (2) Lemma

A graph is feasible and stable if and only if it there is no stable equilibrium where some of the species are extinct, that is also uninvadable on the graph.

Proof: If the graph is feasible and stable, then the interaction matrix for the tree, *α*_*ij*_, is positive definite so the Lyapunov function is concave and the equilibrium is unique; hence there is no other equilibrium in which some species is extinct. If it is not feasible and stable, the existence of a Lyapunov function still implies that there must be some equilibrium, and since the graph is not feasible and stable some species in this equilibrium must be extinct.

## 2. Chains

From Lemma 1 we immediately see that the only allowed subgraph at *α* > 1 is a singlet, as any other subgraph includes a length-2 chain, which is unstable in this range.

In order to calculate the 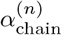, we first derive a rule relating the range of stability and feasibility to the degrees of the vertices of a graph. Using Lemma 2 we show that for any subgraph 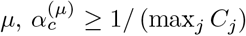, where *C*_*ij*_ is the number of interacting neighbors of species *j*. For a chain, max_*j*_ *C*_*j*_ = 2, and so for any 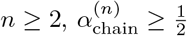.

Indeed, assume that species *j* is extinct at a fixed point. Its growth rate is *g*_*j*_ = 1 – *α*Σ_*k ~ j*_ *N*_*k*_ ≥ 1 − *αC*_*j*_. For *α* < 1/*C*_*j*_ the growth rate would be positive and the equilibrium would be invadable, so in this range species *j* cannot be extinct at an equilibrium. So at *α* < min_*j*_ (1/*C*_*j*_), no species can be extinct at a fixed point, and at the equilibrium all species persist. This behavior is apparent in the histogram of values 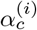 calculated numerically in Fig. 7(A)

## For even length chains 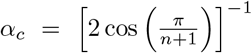

This can be proved by directly checking the feasibility and stability of the chains above 1/2. The interaction matrix representing a chain of length *n* is an *n* × *n* tridiagonal Toeplitz matrix with 1 on the main diagonal and *α* on the diagonals above and below. In this case, the *k*th eigenvalue, with *k* = 1,…, *n*, is

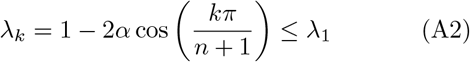

and the chain will be stable for

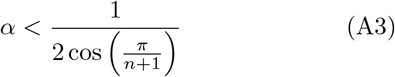

[25]. The chain is also feasible in this range if the abundances are all positive. These abundances equal the sum of the columns of the inverse matrix 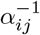. Denoting its components as *σ*_*jk*_, for *j* ≤ *k* they are given by

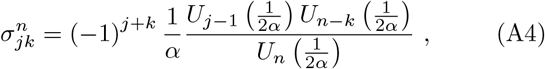

where *U*_*m*_ (*x*) is the *m*-th Chebyshev polynomials of the second kind. The sum over the columns reduces to only two entries of the inverse matrix, so the abundance of the *k*-th species on a chain of length *n* is [36]

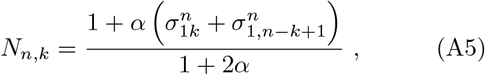

From Lemma 2 we already obtained that 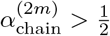, and in the region *α* > 1/2, the *n*th Chebyshev polynomial would be 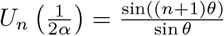, where 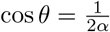. In the region where the chain is stable, 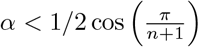, it is straightforward to show that all abundances are positive.

## For odd length chains, *α*_*c*_ = 1/2

For *α* > 1/2, a chain of odd length has an uninvadable equilibrium where species alternate between persistent and extinct, with the persistent species at the odd positions (See Fig 8(A)). The persistent species would have only extinct neighbors and therefore would be stable with abundances *N*_*i*_ = 1. Each extinct species would have two persistent neighbors and its growth rate will be negative, *g* = 1 – 2*α* < 0.

**Figure 8.**
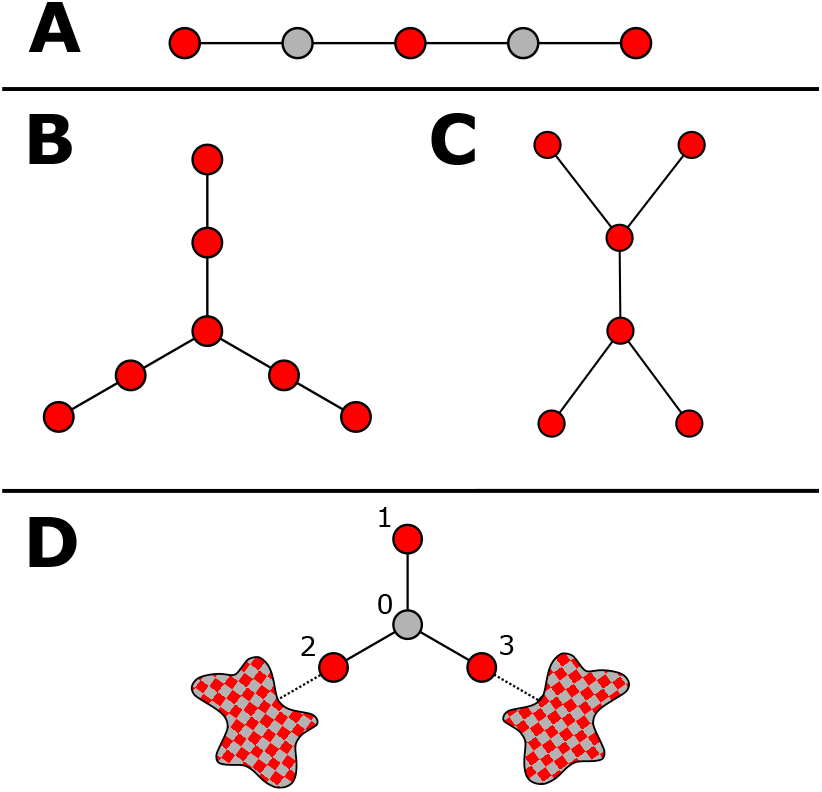
**(A)** The equilibrium at *α* ≥ 1/2 for odd length chains includes alternating persistent (red) and extinct (gray) species, with the persistent species at the odd positions. **(B-C)** Trees which become unstable for *α* > 1/2. All trees that have these as subtrees are also unstable at this range. **(D)** An equilibrium at *α* > 1/2 for all trees that do not have (B-C) as subtrees. Species 0 as defined in the text, at the middle of the junction, is extinct. It breaks the tree into several smaller subtrees of unknown topology, with its neighbors, species 1*,…, m* in the text, persistent leaves of each subtree. Specifically, species 1 has no neighbors besides species 0 and is persistent with *N*_*i*_ = 1.

## 3. Trees that are not chains have *α*_*c*_ ≤ 1/2

To show that such a tree is not allowed above 1/2, we would like to use lemma 2 – if we can remove vertices such that the tree breaks up into chains and the removed vertices are kept from invading by the interactions with the neighbors, we will know that the tree does not also have a fully populated stable fixed point. It is natural to try to remove the species at a junction between three or more branches. We introduce a certain property of junctions and show that if a junction has this property, then when the species at the junction is removed, at equilibrium the abundances of its neighbors are large enough to keep it from invading. If the junction does not have this property, the tree is unstable and therefore if allowed to evolve, some species will naturally become extinct.

## Trees with unstable subtrees

There are two specific trees, shown in Fig. 8(B-C), that are unstable exactly for *α* > 1/2, as can be directly verified. The tree in 8(C) is a subtree of all trees that have two neighboring junctions, and therefore all such trees have *α*_*c*_ ≤ 1/2. Trees that have no neighboring junctions will still have at least one junction (as otherwise they would be chains). For such a tree, if all neighbors of the vertices at the junctions have degree 2, then the tree 8(B) is a subtree and it is also unstable at *α* ≥ 1/2. It remains to be shown that trees that are not chains and do not contain these two subtrees also have *α*_*c*_ ≥ 1/2. These trees must contain at least one junction, as they are not chains; vertices neighboring the junctions can have no more than one other neighbor, otherwise the tree contains the subtree in Fig. 8(C); and at least one neighbor of each junction must have no other neighbors, otherwise the tree contains the subtree in Fig. 8(B). A visualization of such trees, in the case where the junction has 3 neighbors, is shown in Fig. 8(D).

## All trees that are not chains have *α*_*c*_ ≤ 1/2

As we already know that for *α* > 1 all trees are unstable, let us look at trees for some given 1/2 < *α* < 1, and show that they cannot be stable and feasible at this *α*. We will also use the fact that for *α* < 1, the leaves of a tree are always persistent at an equilibrium, as a leaf has only a single neighbor, and therefore if leaf *i* is extinct its growth rate is *g*_*i*_ = 1 – Σ_*j*_ *α*_*ij*_*N*_*j*_ ≥ 1 – *α* > 0. Further, its abundance is bounded from below, *N*_*i*_ (*α*) = 1 – *α* Σ_*j*~*i*_ *N*_*j*_ ≥ 1 – *α*.

If a tree has either of the trees in Fig 8(B-C) as sub-trees, then we already showed that it is unstable at *α*. Otherwise, as explained at the end of the previous part, this tree must have a junction with no neighboring junctions, and at least one neighbor with degree 1, as shown in Fig 8(D). We now find an equilibrium with extinct species for trees with such a topology.

Mark the species in the middle of the junction as species 0, and its neighbor that has degree 1 as species 1. Species 0 has additional neighbors 2,…, *m*, with *m* ≥ 3, as indicated in Fig 8(D). Each of these species has at most one more neighbor besides species 0, otherwise there would be a neighboring junction to species 0.

We now show that an equilibrium exists where species 0 is extinct. As species 0 goes extinct, the tree separates into *m* distinct subtrees of size < *N*. Let us examine the equilibria of these subtrees. The first subtree includes only species 1 (as it had no other neighbor besides species 0), so it has an equilibrium where species 1 is persistent with *N*_1_ = 1. All other subtrees also have an equilibrium, because of the existence of a Lyapunov function for each subtree separately. We now need only check that species 0 cannot invade at this equilibrium.

Specifically, as species 2,…, *m* are leaves of their respective trees, they must be persistent at these equilibria, and have abundances *N*_*i*_ (*α*) ≥ 1 – *α*. The growth rate of species 0 is therefore

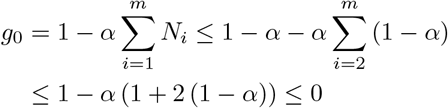

and indeed it cannot invade. So the tree has an equilibrium with extinct species, and thus it cannot be stable and feasible.

## Appendix B Effects of invadability and non-tree subgraphs

In the main text, we explain that the jumps in the relative diversity *ϕ* (*α*) in the region *α* > 1/2 result from changes in the stability and feasibility of trees. We neglect the effect of changes in the feasibility and stability of subgraphs that are not trees, and the invadability of extinct species. In this section we will discuss these assumptions and show that such changes do not appear to generate additional jumps in *ϕ*.

## 1. Invadability

In this section, we will show that for *α* > 1/2, most extinct species would not change their invadability within the ranges where there is no change in feasibility and stability of trees. In the cases where they do, the jumps generated, if they exist, are so small that we do not observe them in our simulations

A change in invadability occurs at *α*-values where there is a sign change of the growth rate of an extinct species, *g*_*i*_ (*α*) = 1 – *α*Σ_*j*~*i*_ *N*_*j*_ (*α*). For *α* > 1/2 the only allowed trees are even length chains, with a finite number of possible abundances *N*_*j*_ (*α*), and so a finite number of possible growth rates *g*_*i*_ (*α*), depending on the different possible combinations of neighboring species. For example, in the range where the only allowed trees are singlets and length 2 chains, the only possible abundances are 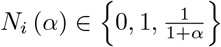 and for a *C*-regular graph, this gives 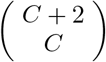 possible growth rates. We checked for such sign changes for *C* = 3 in the range that allows chains of up to length 2,4,6 and 8, about *α* ∈ [0.52, 1]. We exclude the values 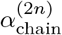 where we already know there are jumps in *ϕ* due to changes in chain stability. We stop at the maximal chain length of 8 as the number of possible growth rates grows very fast with the maximal allowed chain length. In the ranges allowing chains of up to length 2 and 4, no growth rate changes sign; in the range allowing chains of up to length 6, one possible growth rate out of 120 changes sign; in the range allowing chains of up to length 8, 5 out of 364 possible growth rates change sign.

In Fig. 9(B), we show *ϕ* (*α*) around a value of *α* where one of these changes in growth rates occur, along with the specific combination of neighbors that generates the change. Even with an increase in the pool size *S*, we see no jump in the value of *ϕ* within statistical error. These results could be expected, as each growth rate occurs only when an extinct species has neighboring chains of very specific lengths, which happens very infrequently. However, this does not mean that invadability is unimportant, as it drives the changes in the allowed subgraphs. For example, chains of length 4 become allowed at the value of *α* such that an extinct species which neighbors a length two chain (of persistent species) and another single species can invade, so that all the sites stick together as a length 4 chain. This is just another way of describing the result above, that a graph is not allowed if removing some species from it leads to a subgraph such that the removed species cannot invade.

**Figure 9.**
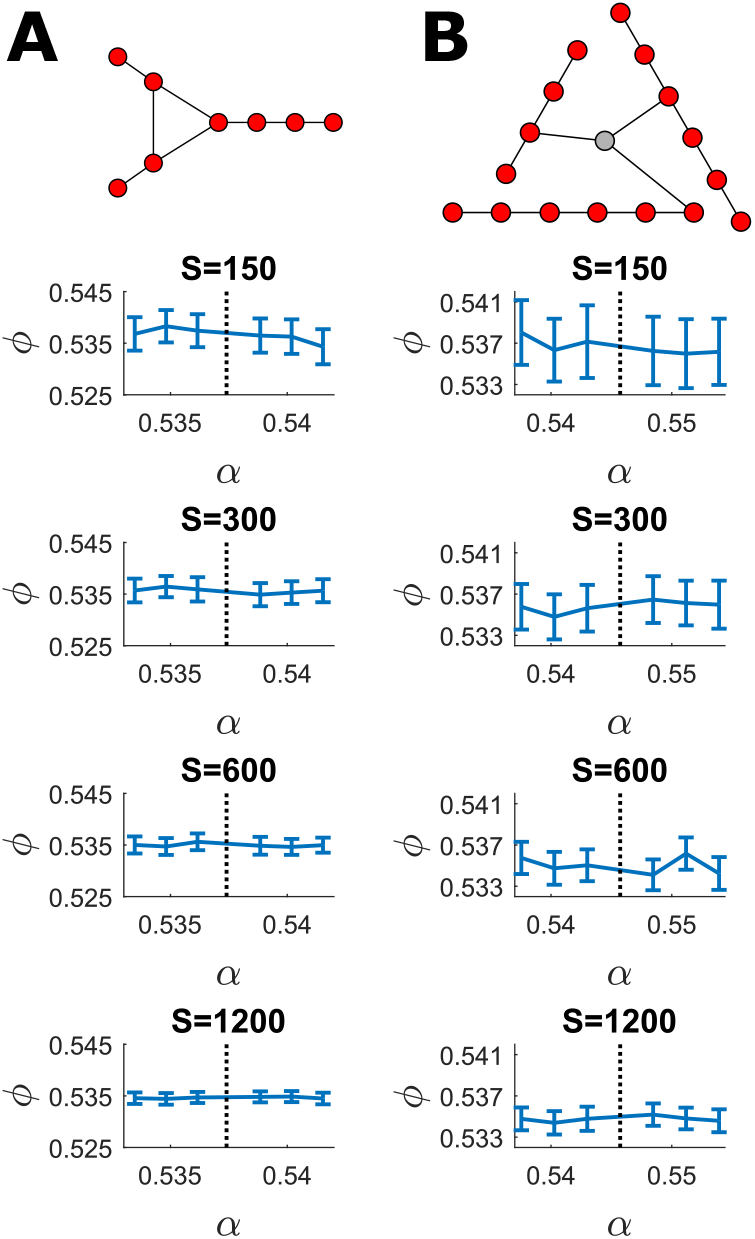
Non-tree subgraphs and invadability changes do not generate jumps in *ϕ* **at** *α* > 1/2. The dependence of relative diversity *ϕ* on the interaction strength *α* for increasing pool sizes *S*, in two cases **(A)** Around *α* ≈ 0.537, the value at which a non-tree subgraph becomes allowed. The subgraph is shown at the top **(B)** Around *α* ≈ 0.546, the value where an extinct species (in gray) with a set of persistent neighbors as shown will change the sign of its growth rate, and so its invadability.

## 2. Non-tree subgraphs

As mentioned in the main text, as sparse graphs are tree-like and short cycles are rare, we expect to see no jumps generated by subgraphs that are not trees. Fig. 9(A) shows an example for a specific subgraph that includes a cycle: *ϕ* (*α*) displays no jump around the critical value where this subgraph becomes allowed, even as we increase *S*.

## Appendix C Collective transitions with heterogeneity

In this section we continue to examine the two collective transitions, the transition from multiple to unique equilibria and the percolation transition, in cases where interaction strengths and vertex degrees are not constant across the network. The behavior at the transitions is shown in Fig. (4) in the main text for heterogeneous interaction strengths, and here in the top panels of Fig (10) for variability in vertex degree modeled by an Erdős-Rényi graph. For both cases, the transition from multiple to unique equilibria becomes sharper as *S* increases (within the range checked numerically), with the probability of a unique equilibrium approaching a step function. The percolation transition in both cases is qualitatively similar to the transition that occurs in the case with no heterogeneity, as well as to standard site percolation, see Section D.

**Figure 10.**
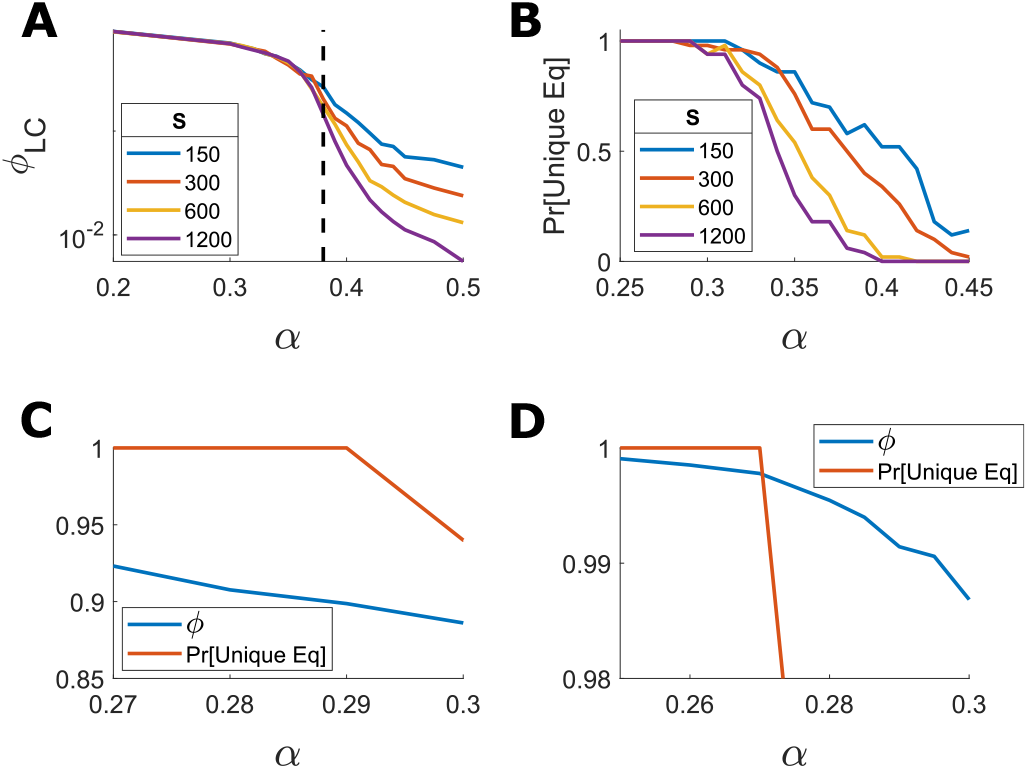
Collective transitions with heterogeneity. Results in panels (A-B) are the equivalent of Fig. 3(C,D), here for heterogeneity in degree the transitions. As in Fig. 3(C,D), the transitions become sharper as *S* increases, within the range of *S* tested numerically. They show simulation results for Erdős-Rényi graphs with average degree *C* = 3 and several values of *S*. **(A)** Percolation transition: The fraction of species in the largest connected component as a function of *α*. At low values of *α* a finite fraction of species belong to the largest component, and above a transition the fraction of species decreases with *S*. **(B)** Multiple to unique equilibrium transition: The probability of having a unique equilibrium as a function of *α*. **(C-D)** For both types of heterogeneity some species go extinct (a loss of feasibility of the entire system) before loss of stability: The fraction of surviving species *ϕ* drops below 1 in the unique equilibrium phase. Results are shown for *S* = 1200, *C* = 3. The probability for a unique equilibrium is shown in red, and *ϕ* in blue. **(C)** Heterogeneity in degree, Erdős-Rényi graphs **(D)** Heterogeneity in interaction strength, *σ* = 0.1.

Fig. 10(C,D) show that for both types of heterogeneity, *ϕ* drops below 1 before the transition to multiple equilibria, for *α* < *α*_UE_. Therefore, the feasibility of the entire system is lost before its stability. For heterogeneous interaction strengths, this follows from Lemma (3) in Appendix E.

## Appendix D Comparison to standard percolation

Here we elaborate on the comparison in the main text between percolation in our model and standard site percolation. In standard percolation, each vertex is taken to be “present” with a given probability *p*, and for *C*-regular graphs the percolation transition is known to occur at 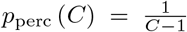 [31]. Fig. 11 compares three cases: standard percolation on a random regular graph, percolation in the equal-*α* model where interaction strength and degree are constant, and for heterogeneous interaction strengths. For each we show the dependence of the fraction of species in the largest connected component, *ϕ*_*LC*_, on the fraction of surviving species, *ϕ*, or on *p* for standard percolation. As mentioned in the main text, in all cases the behavior close to the transition is qualitatively similar. We use this similarity to estimate *α*_perc_ in our model, as the value of *α* where the fraction of species in the largest component grows as *S*^−1/3^, as in known to occur at p_perc_ for standard site percolation [31]. As mentioned, for our model *ϕ*_perc_ > 1/2 = *p*_perc_, due to the fact that persistent species are anticorrelated, tending not to be adjacent to one another.

**Figure 11.**
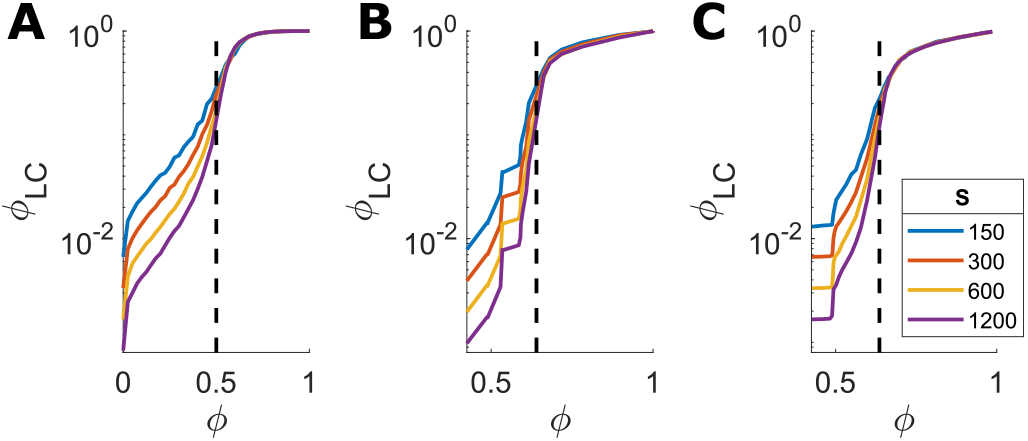
Percolation in our model is qualitatively similar to standard site-percolation, near the transition. The dependence of the fraction of species in the largest connected component on the fraction of all persistent species *ϕ*, for *C* = 3 and several pool sizes *S*. **(A)** Standard site-percolation: vertices are taken to be present with probability *ϕ*, independently for each vertex. The percolation transition occurs at *ϕ*_perc_ = 1/2, where *ϕ*_LC_ ~ *S*^−1/3^. **(B)** The equal-*α* model, with constant interaction strength and vertex degree, *ϕ*_perc_ ≈ 0.64. **(C)** The model with variability *σ* = 0.1 in interaction strength, *ϕ*_perc_ ≈ 0.636.

## Appendix E Subgraph emergence rule

To prove the result quoted in the main text, we first prove a Lemma, which is interesting in its own right. We use the term “generically” for “with probability approaching one for large numbers of species”.

### (3) Lemma

Consider a system with symmetric (*α*_*ij*_ = *α*_*ji*_) and competitive (*α*_*ij*_ ≥ 0) interactions, sampled from some continuous distribution (such as a Gaussian distribution as in the main text). Suppose that the *α*_*ij*_ are changed continuously by shifting *m* ≡ mean (*α*_*ij*_) (other continuous shifts are also possible). Assume that the graph is feasible and stable in some range below *m* = *α*_*c*_, and not in a range above it. Then generically, it is feasibility that breaks at *α*_*c*_, by a single species’ abundance going to zero, while stability continues to hold.

Proof: We prove this by contradiction. Assume to the contrary that at *m* = *α*_*c*_ the graph becomes unstable. As the matrix *α* is symmetric it can be diagonalized. Let 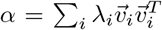 be its eigen-decomposition, where *T* denotes the transpose operation, 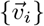 are column eigenvectors, and *λ*_*i*_ the corresponding eigenvalues, with *λ*_1_ < *λ*_2_ <… (generically there is no degeneracy). Note that the values of quantities in this decomposition depend on m. These values are equilibrium solutions to Eq. (1); fromfeasibility up to *α*_*c*_, all *N*_*i*_ > 0 so,

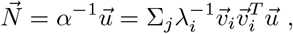

where 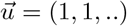. By assumption, the system becomes unstable at 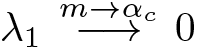. Since generically 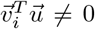, the first term dominates near *α*_*c*_,

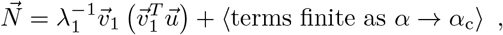

so 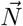 diverges at *m* → *α*_*c*_. Using *α*_*ij*_ ≥ 0 and feasibility, *N*_*i*_ = 1 – Σ_*j*_ *α*_*ij*_*N*_*j*_ ≤ 1. Therefore, the divergence of the values of 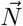 must be towards −∞, and so the *N*_*i*_-values must cross zero at m *smaller than α*_*c*_, in contradiction to the assumption. QED

Applying this lemma, a subgraph that loses feasibility at *α*_*c*_ generically does so by only one species having N_*i*_ → 0. The remaining graph is still feasible and stable at *α*_*c*_ and for at least some range [*α*_*c*_, *α*_*c*_ + *ε*] above it (because the stability and abundances of the remaining species change continuously). In the case of trees, removing a vertex splits the tree into multiple trees, see Fig. 5.

Without heterogeneity (when all *α*_*ij*_ = *α*) all trees have *α*_*c*_ ≤ 1/2, so it is interesting to consider the case where all the *α*_*ij*_ connecting to the extinct species *N*_*i*_ satisfy *α*_*ij*_ < 1/2. In this case, the extinct species has 0 = *N*_*i*_ = 1 – Σ_*j*_ *α*_*ij*_*N*_*j*_ > 1 – *C*/2, where *C* is the degree of species *i*, so *C* > 2 and the tree will split into at least three parts.

## Notes

### Competing Interest Statement

The authors have declared no competing interest.

